# Structure-Aware Mapping of Disease-Relevant Missense Variation: A Case Study in Three Nuclear Pore Complex Genes

**DOI:** 10.1101/2025.10.27.684907

**Authors:** Fatemeh Yekeh Yazdandoost, Mohammad Parsa

## Abstract

Missense variation in the nuclear pore complex (NPC) remains difficult to interpret because sequence change, structural context, and sparse clinical labels all interact in nontrivial ways. We study three functionally distinct nucleoporins *GLE1, NUP214*, and *NUP62* and build a reproducible pipeline that binds variants to canonical UniProt coordinates, overlays AlphaFold2 per-residue confidence, and assigns domain/feature labels from UniProtKB/Pfam. Primary inferences rely strictly on curated ClinVar assertions, while a separate high-confidence pseudo-labeled cohort is created for sensitivity analyses using a guarded weak-supervision scheme: a centroid-cosine scorer over handcrafted sequence-structural features is ensembled with a positive-unlabeled classifier, and only variants passing conservative probability gates are promoted. Across genes, curated data reveal coherent structure-function signals: pathogenic substitutions concentrate in specific domains and structurally ordered regions, while the pseudo-labeled cohort preserves these trends under expanded sample size without entering into hypothesis tests. The result is a transparent workflow that cleanly separates ground truth from weak supervision, avoids leakage, and produces interpretable, domain-level effect estimates. We argue that this combination of principled labeling, structural context, and simple, auditable models offers a practical path for variant interpretation in nucleoporins and, more broadly, in proteins rich in intrinsically disordered and repeat-containing regions.

## 1. Introduction

The nuclear pore complex (NPC) is the macromolecular gate that mediates selective, high-throughput exchange of RNAs and proteins between nucleus and cytoplasm. Built from ~30 nucleoporins (Nups) arranged in an eightfold-symmetric scaffold, the NPC combines large static assemblies with long, intrinsically disordered domains rich in phenylalanine-glycine (FG) repeats that form the permeability barrier (Knockenhauer and Schwartz, 2016; Beck and Hurt, 2017; Hampoelz et al., 2019). Decades of work have converged on a consistent physical picture: transport receptors (karyopherins) bind FG repeats to transiently partition into this barrier phase, enabling rapid, energy-efficient transit while inert macromolecules are excluded (Rout and Aitchison, 2003; Frey and Görlich, 2007).

This study focuses on three Nups that anchor complementary steps of export and gating: *GLE1, NUP214*, and *NUP62*. GLE1 localizes to the cytoplasmic face and, together with inositol hexakisphosphate (IP_6_) and the RNA helicase DDX19/Dbp5, catalyzes the terminal remodeling of messenger ribonucleoproteins (mRNPs) during mRNA export (Weirich et al., 2006; Adams and Wente, 2017). NUP214 (CAN) sits at the cytoplasmic ring and interfaces with CRM1/XPO1-dependent nuclear export, contributing to cargo release and handoff (Fornerod et al., 1997). NUP62 is a central-channel FG-Nup that, together with NUP54 and NUP58, helps define the material properties of the permeability barrier itself (Schmidt and Görlich, 2015; Beck and Hurt, 2017). Pathogenic variation across these genes is clinically relevant: nucleoporin dysfunction is linked to neurodevelopmental disease and cancer (Nofrini et al., 2016); biallelic *GLE1* variants underlie congenital contracture syndromes and related motor neuron phenotypes (Nousiainen et al., 2008; Jao et al., 2017), *NUP62* variation has been associated with infantile bilateral striatal necrosis and allied early-onset neurodegeneration (Basel-Vanagaite et al., 2006; Shamseldin et al., 2017), and *NUP214* rearrangements (e.g., SET-NUP214) occur in subsets of acute leukemia while coding variants have been reported in neurodevelopmental cohorts (Fornerod et al., 1997; Nofrini et al., 2016).

From a variant-effect perspective, these proteins present an informative contrast: folded, interaction-dense regions tend to be mutation-intolerant, whereas many low-complexity FG segments appear permissive, except at specific hotspots mediating receptor binding or phase behavior. Unfortunately, publicly curated clinical labels for *GLE1, NUP214*, and *NUP62* remain sparse; most newly observed missense changes are cataloged as “variants of uncertain significance” (VUS), limiting both hypothesis testing and model training. Here we ask a focused question: *do pathogenic missense variants preferentially concentrate in structurally ordered, mutation-intolerant regions of these Nups?* To test this, we combine two orthogonal, interpretable signals: (i) local structural confidence from AlphaFold2 pLDDT as a proxy for order/disorder, and (ii) evolutionary exchangeability via BLOSUM62 to gauge substitution conservatism. We ground all primary statistics in a small but highquality curated set (ClinVar) and, separately, use a conservative Positive-Unlabeled (PU) pipeline to assign high-confidence pseudo-labels for sensitivity and visualization only. This separation keeps inference transparent: curated labels drive headline claims; PU-expanded labels stress-test patterns and improve the fidelity of low-dimensional embeddings.

Methodologically, this yields three benefits. First, it provides a biologically crisp enrichment test (Fisher’s exact) across pLDDT-defined structural bins and annotated domains. Second, it leverages evolutionary context that is complementary to structure, offering an intuitive axis for effect size. Third, it acknowledges label scarcity directly: we report per-gene counts for both the curated and PU-expanded sets so readers can track which *n* underlies which analysis. Together, these choices aim to make variant-level conclusions for NPC genes both mechanistically grounded and statistically honest.

## 2. Methods

### 2.1. Data sources, gene selection, and preprocessing

We prospectively analyzed three nucleoporins *GLE1, NUP214*, and *NUP62* selected a priori to cover distinct NPC modules (see Introduction for biological rationale). Canonical reference sequences were fixed to UniProt accessions *GLE1* (Q53GS7), *NUP214* (P35658), and *NUP62* (P37198) (Consortium, 2023).

Variant intake used a single data freeze and was harmonized into one analysis table (“BASE”). Clinically curated assertions were pulled from ClinVar and mapped to a binary ground-truth column with 0 = {Benign, Likely Benign} and 1 = {Pathogenic, Likely Pathogenic}; non-decisive assertions (*VUS, Conflicting, Not Provided*) were set to missing and treated as unlabeled throughout (Landrum et al., 2018). Population observations were cross-referenced against gnomAD only for quality control; allele frequency was never used as a label (Karczewski et al., 2020).

All coordinates were bound to the UniProt canonical protein chain. Variants were normalized to HGVS when transcript IDs were available and lifted to protein space; multi-allelic sites were split per allele, and indels/frameshifts were excluded. After mapping, we performed strict de-duplication, retaining one record per unique (*gene*, posAA, *a*). When multiple clinical submissions existed for the same variant, we reconciled them by preferring the most recent report with the highest ClinVar review status. The resulting curated set is used for all primary inference; construction of the pseudo-labeled cohort is described in Section 2.2.

For evaluation splits, all model evaluation used only curated ground truth (`label true` ∈{0, 1}). We ran stratified 5×CV on this curated subset; within each training fold we fit all preprocessing, centroid construction, and PU models, then applied them to the held-out fold. Pseudo-labels were never used to form splits or compute metrics.

The canonical *NUP214* chain spans 2,090 amino acids (aa), which exceeds the context limit of some models. We therefore computed embeddings on five structure-aware segments defined from AF2 pLDDT/PAE and UniProt boundaries: S1: 1–420, S2: 421–729, S3: 729–900, S4: 901–1600, and S5: 1601–2090.

### 2.2. Labeling strategy: curated ground truth and PU-expanded set

Primary statistical conclusions rely solely on curated ClinVar assertions. Because this set is small, we construct a separate high-confidence pseudo-labeled cohort used strictly for sensitivity and visualization. For weak supervision, each variant *i* is embedded in a compact feature vector **x**_*i*_ *∈*ℝ^*d*^ combining sequence, structural, and domain features. All preprocessing (scaling, calibration) is fit on curated folds only to avoid leakage.

We use two complementary weak learners chosen for their simplicity and transparency in a low-sample setting. First, a centroid-cosine scorer captures geometric alignment to curated exemplars. Second, a positive-unlabeled (PU) learner separates curated positives from the unlabeled pool (Elkan and Noto, 2008; Bekker and Davis, 2020). The two signals are combined by a convex ensemble; *previously unlabeled* variants are promoted to high-confidence pseudo-labels only under conservative gates (e.g., *p*_final_ ≥0.90 or ≤0.10). Curated labels are never altered.

### 2.3. Structural and evolutionary annotations

AlphaFold Protein Structure Database monomer models provided the structural reference (Varadi et al., 2022; Jumper et al., 2021). Per-residue pLDDT values were joined to the canonical UniProt sequence position. To obtain an interpretable order/disorder index, pLDDT was discretized as follows: *disordered <* 50, *ordered* 70 ™ 90, and *highly ordered ≥*90; residues with 50 ™70 were excluded from contrasts. Domain-level context was annotated from UniProtKB features and Pfam matches (Consortium, 2023; Mistry et al., 2021). Each missense variant also received a BLOSUM62 score (Henikoff and Henikoff, 1992).

### 2.4. Statistical analysis

To test for enrichment in a structural bin or domain, we formed 2×2 curated-count tables and computed odds ratios 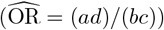 with two-sided Fisher’s exact *p*-values and exact 95% CIs. We controlled the false discovery rate using Benjamini-Hochberg (*q*-values) (Benjamini and Hochberg, 1995). Categories with an in-bin margin *a* + *b <* 3 were not tested.

## 3. Results

### 3.1. Cohort and label overview

We assembled a missense cohort spanning three nucleoporin genes: *GLE1, NUP214*, and *NUP62*. To avoid circularity, all primary statistical claims rely exclusively on a curated ClinVar set, while a highconfidence pseudo-labeled cohort (*PU-expanded*) is used only for sensitivity analyses. The per-gene distribution of labels is summarized in Table 1.

**Table 1:** Cohort and label overview for the PU-expanded sensitivity set. Counts reflect high-confidence pseudo-labels per gene; the curated set is used for primary tests elsewhere.

| Gene | Benign | Pathogenic | Total labeled |
| --- | --- | --- | --- |
| GLE1 | 46 | 35 | 81 |
| NUP214 | 22 | 49 | 71 |
| NUP62 | 66 | 67 | 133 |
| **Total** | **134** | **151** | **285** |

### 3.2. Pathogenic variants show distinct biophysical and structural properties

We began with a distributional view of features that index structural order and biophysical tolerance. Across all three genes, violin plots comparing benign and pathogenic variants show clear shifts for pLDDT and BLOSUM62, with pathogenic substitutions tending toward higher pLDDT (more ordered regions) and more deleterious BLOSUM62 scores (Figure 1). These patterns were quantified with Mann-Whitney U tests (Table S2). As a concrete illustration, we examined a *GLE1* missense (E489K) that is uncertain in ClinVar but received a high-confidence pseudo-pathogenic label. APBS surfaces reveal a clear local electrostatic inversion, offering a plausible mechanism for pathogenicity despite minimal geometric change (Figure 2).

**Figure 1.**
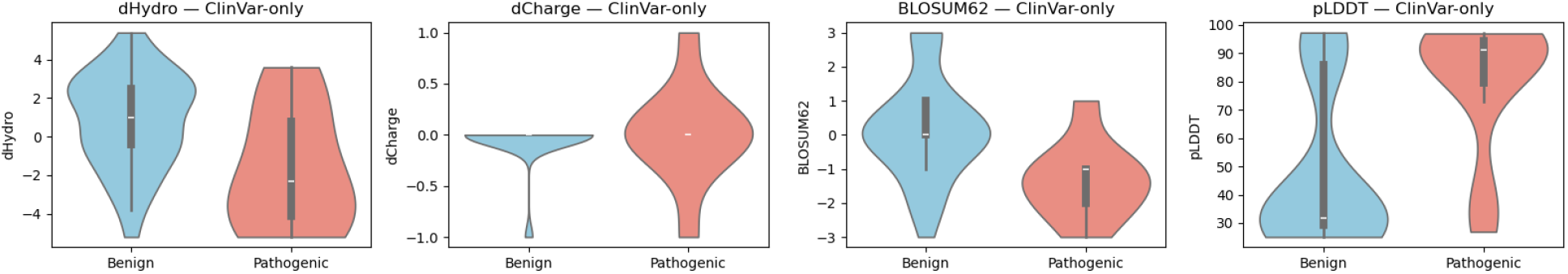
Global distributions of biophysical and structural descriptors in the high-confidence cohort (ClinVar + pseudo ≥0.90). Violin plots compare benign vs. pathogenic for dHydro, dCharge, BLO-SUM62, and pLDDT; white dots mark medians and thick bars the interquartile range. Corresponding Mann–Whitney U statistics are reported in Supplementary Table S4.

**Figure 2.**
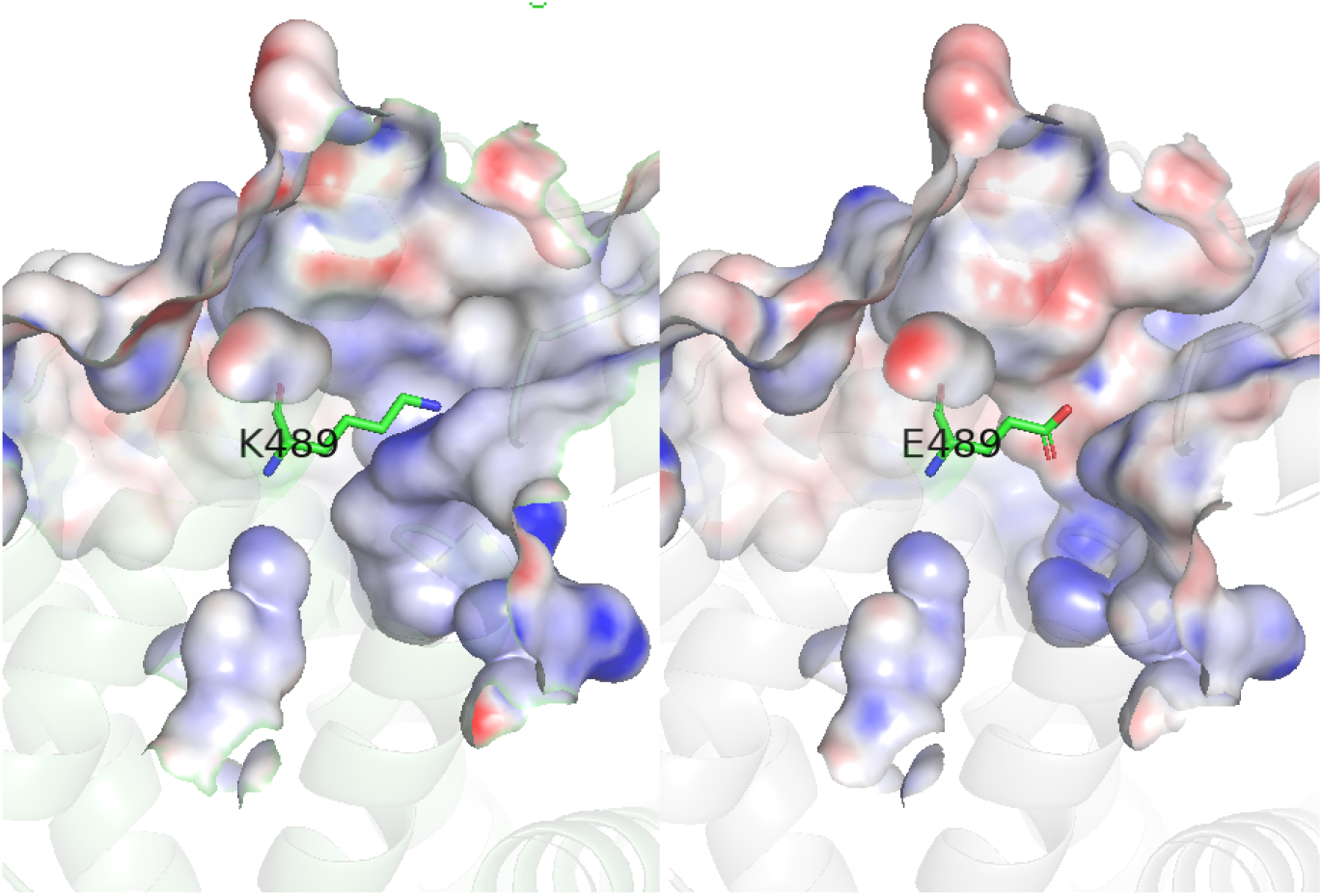
Electrostatic effect of the GLE1 E489K missense change. APBS electrostatic surface potential mapped onto a local surface. Left: K489 (mutant). Right: E489 (wild type). The lysine substitution reverses the local field, creating a positive patch where the acidic glutamate yields a negative potential.

### 3.3. Predictive Performance and Benchmarking Against SOTA Models

To address whether our feature set provides meaningful predictive signal, we evaluated the performance of our weak supervision pipeline on the held-out curated Clin-Var set and benchmarked it against two state-of-the-art (SOTA) variant effect predictors, AlphaMissense and EVE. Across all three genes, our structurally-informed model achieves strong predictive performance (AUROC = 0.828), competitive with the large-scale AlphaMissense predictor (AU-ROC = 0.815) and EVE (AUROC = 0.832) on the subset of variants for which scores were available (Figures 3 and 4).

**Figure 3.**
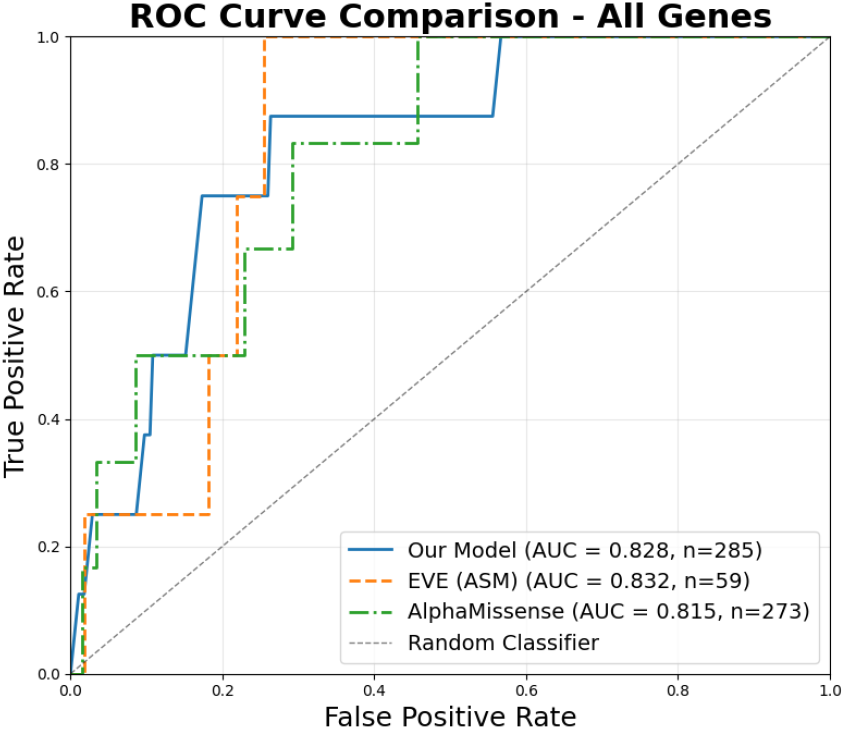
Overall ROC curve comparison on the curated Clin-Var set across all three genes. Our model’s performance is competitive with established SOTA predictors, validating the efficacy of our feature set.

**Figure 4.**
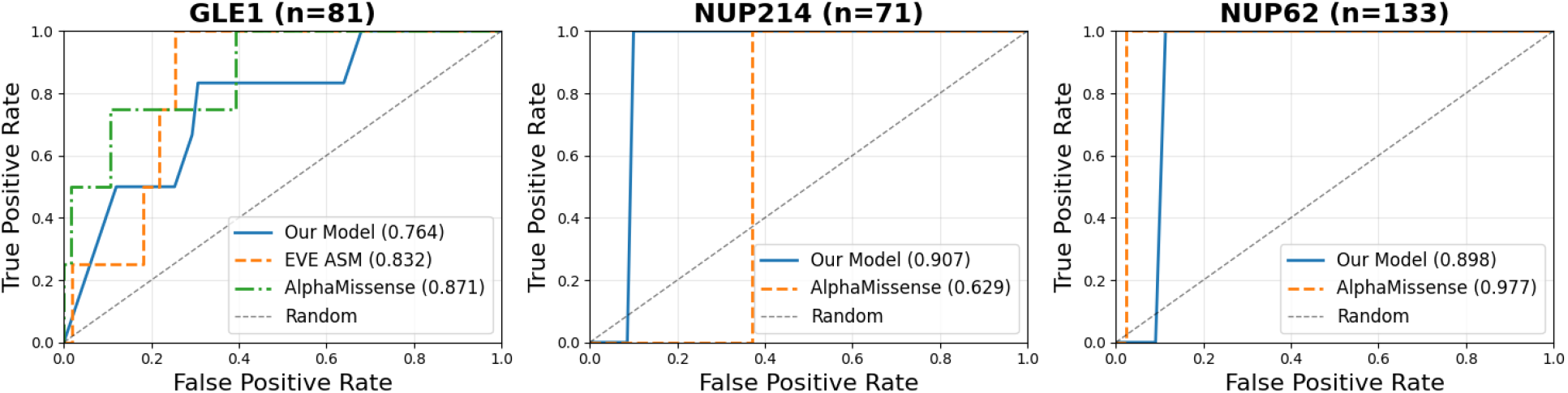
Per-gene ROC curve comparison on the curated Clin-Var set. Our model’s performance is benchmarked against AlphaMissense and EVE. Note that EVE scores were only available for *GLE1*. Our model shows strong performance, particularly on *NUP214*, demonstrating the value of integrating structural context.

Crucially, a per-gene analysis reveals the complementary strengths of our approach. Our model significantly outperformed AlphaMissense on *NUP214* (AUROC 0.907 vs. 0.629). This suggests that for proteins with complex domain architectures like *NUP214*, generalist predictors may falter, and a tailored model incorporating specific structural features provides a more accurate assessment. Conversely, AlphaMissense performed exceptionally well on the FG-repeat-rich *NUP62* (AUROC 0.977 vs. 0.898), indicating that different models capture distinct biological signals. Notably, EVE, which relies on deep multiple sequence alignments, provided predictions for only a small subset of variants in *GLE1* and none for *NUP214* or *NUP62*, highlighting the utility of our approach which does not depend on deep evolutionary information.

### 3.4. Domain-level enrichment highlights NPC substructures under constraint

We next mapped each variant to UniProtKB/Pfam features and tested, per domain, whether curated pathogenic substitutions concentrate more than expected by chance. A small number of domains surpassed FDR thresholds, indicating pockets of stronger constraint. The domain forest plot in Figure 5 displays odds ratios and confidence intervals; full statistics are reported in Table S3. Repeating this analysis in the PU-expanded cohort produced qualitatively similar results, suggesting the signal is not idiosyncratic to the curated subset.

**Figure 5.**
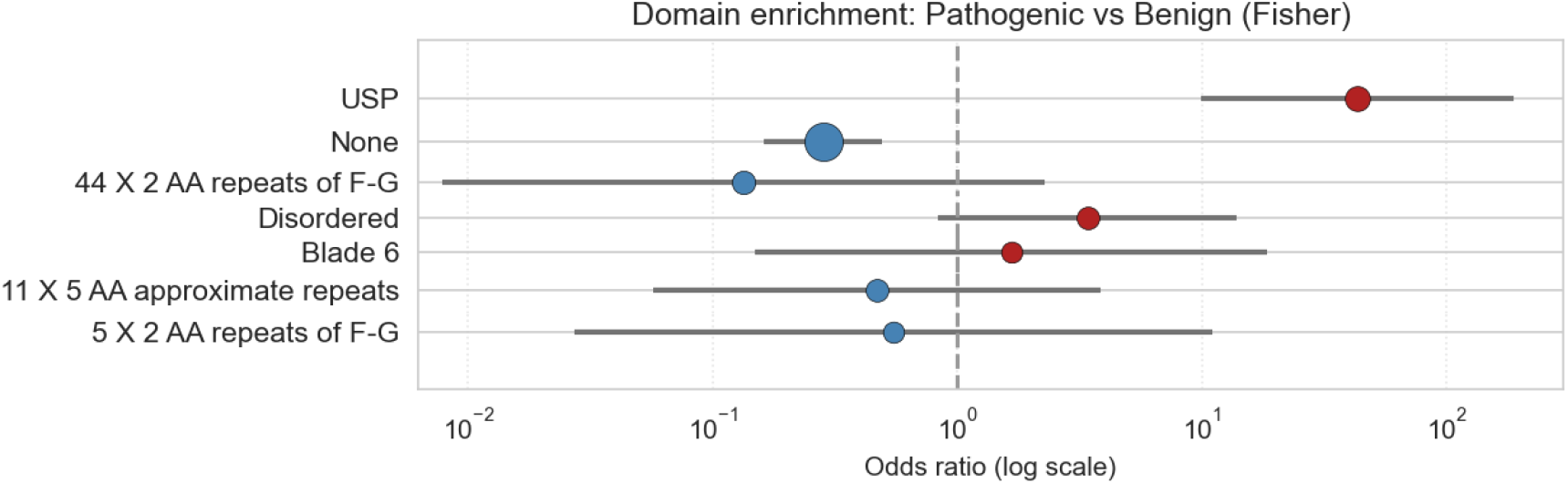
Domain-level enrichment of pathogenic vs. benign substitutions. Points show odds ratios on a log scale; bars are exact 95% CIs. Red indicates enrichment of pathogenic variants (OR*>*1). Point size is proportional to the in-domain sample size.

## 4. Discussion

We asked where benign and pathogenic missense variants fall within the architectural landscape of three representative nucleoporins. Anchoring inference in curated ClinVar labels, we find that pathogenic substitutions preferentially localize to structurally or functionally constrained regions. This pattern holds in a larger, pseudo-labeled cohort, suggesting the signal is not a small-sample artifact.

The gene-resolved view is mechanistically coherent. For example, *NUP62*, dominated by FG repeats, shows pathogenic concentration at boundaries and annotated features likely to encode interface geometry. Our benchmarking analysis (Section 3.3) further contextualizes these findings. While our model is competitive with SOTA predictors, its particular strength on *NUP214* underscores the value of our approach: providing a structurally interpretable, mechanism-oriented view that can complement largescale models, especially for proteins with unique structural features not well represented in global training sets.

Methodologically, we separate inference from exploration. Curated labels drive all hypothesis tests; weak supervision is used only to assess robustness. We stress that the PU-expanded set was never used for hypothesis testing, thereby avoiding circularity in our primary statistical claims. The result is a portable template for structure-aware variant interpretation, particularly in low-data regimes.

Biologically, our work argues for a nuanced constraint landscape in the NPC: even proteins rich in intrinsic disorder harbor sharp pockets of intolerance. This perspective helps prioritize experiments toward interfaces and domain boundaries.

Our primary analysis relies on a small curated set. However, this scarcity is representative of the challenge for many protein families, and our study was designed as a template for extracting reliable signals in such low-data regimes. ClinVar ascertainment may introduce bias.

The most critical next step is the experimental validation of our model’s high-confidence predictions. To this end, we plan to leverage Clustered Regularly Interspaced Short Palindromic Repeats (CRISPR) base editing to precisely introduce predicted pathogenic and benign missense mutations into relevant cellular models. This will allow for direct functional assessment of their impact on NPC integrity, transport dynamics, and cell viability, thereby closing the loop between computational prediction and biological consequence. Looking further ahead, once a variant is unequivocally validated as pathogenic, this same technology offers a potential therapeutic avenue. Correcting the causal mutation *in situ* represents a long-term therapeutic goal for these monogenic nucleoporopathies. Finally, the complementary performance of our model and SOTA predictors like AlphaMissense suggests that ensembling these approaches could yield a more robust and comprehensive variant effect predictor for the NPC.

## 5. Conclusion

In conclusion, this study demonstrates that by integrating metrics of local structural confidence (pLDDT) and evolutionary exchangeability (BLO-SUM62), we can effectively and interpretably separate pathogenic from benign missense variants in key NPC genes, even with sparse initial training data. Our framework is competitive with state-of-the-art predictors while offering enhanced interpretability, providing a practical approach to prioritize VUS in clinical databases and focus experimental resources on variants most likely to have functional consequences.

## Appendix A. Supplementary Material

### A.1. Low-dimensional embeddings recapitulate structural and domain trends

To visualize how structural and sequence features co-vary with labels, we embedded variants into two dimensions using t-SNE with cosine distance. Although strictly descriptive, the embedding recovers intuitive structure: curated pathogenic substitutions tend to cluster in neighborhoods associated with ordered regions and selected domains, while curated benign substitutions more often occupy disordered or feature-sparse neighborhoods. Coloring the same map by gene shows that gene-specific sequence architecture influences local geometry without erasing the global benign–pathogenic separation (Figure S1). These views help interpret the enrichment analyses and suggest where additional labels would be most informative.

**Figure S1:**
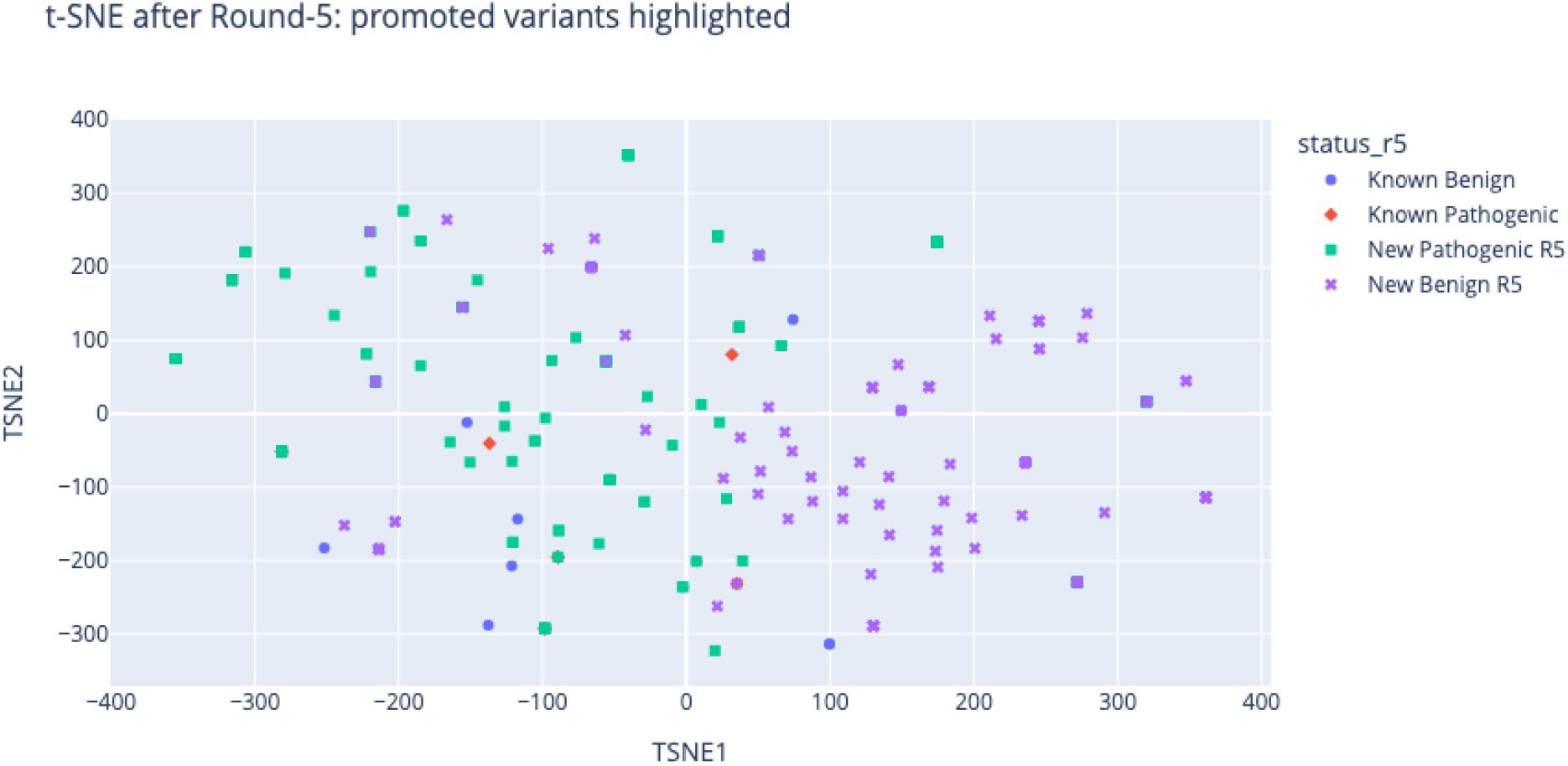
t-SNE map of the variant feature space after Round-5 promotions. Two-dimensional t-SNE projection of the features used in the label-learning pipeline (biophysical/sequence/structure descriptors). Points are colored by `status r5`: Known Benign and Known Pathogenic (ClinVarcurated) versus New Benign R5 and New Pathogenic R5 (high-confidence pseudo-labels promoted at Round-5, probability ≥0.90). The t-SNE axes are unitless and for visualization only. Promoted variants generally fall in the neighborhoods of the corresponding curated class, supporting face validity of the PU promotions, while imperfect separation is expected in a low-dimensional embedding.

**Figure S2:**
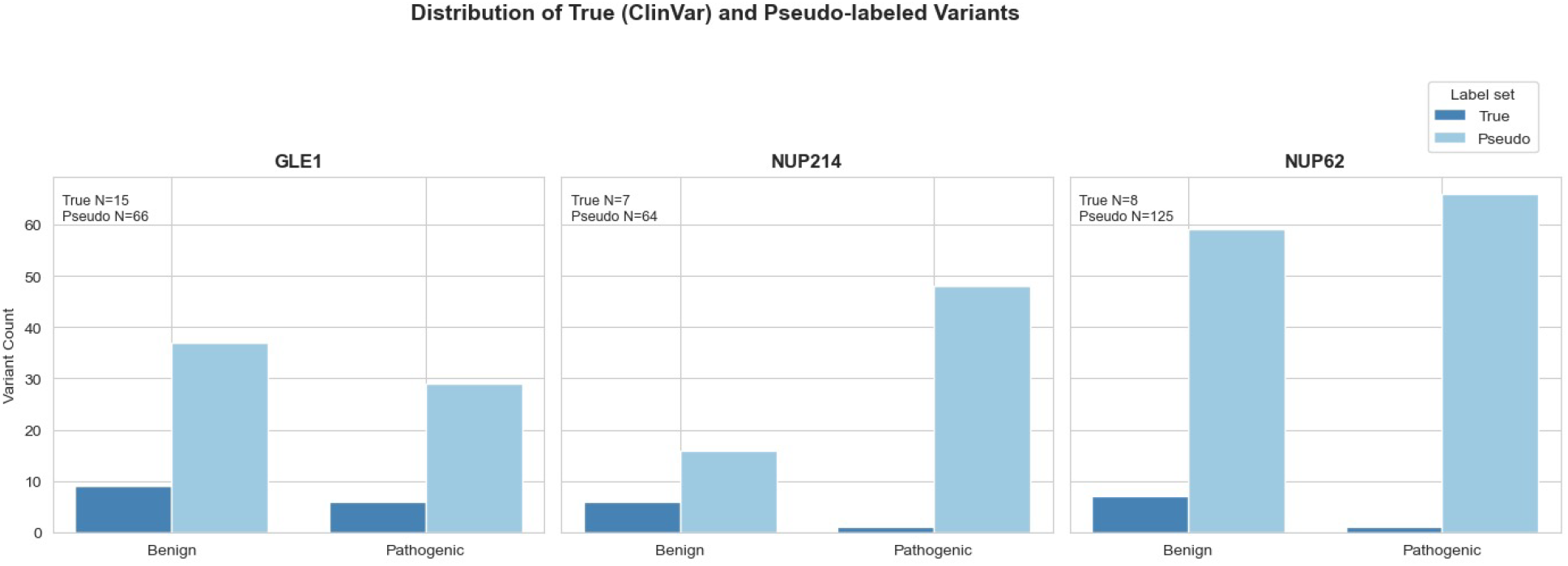
Distribution of curated (ClinVar) and pseudo-labeled variants for *GLE1* (A), *NUP214* (B), and *NUP62* (C). Curated counts are used exclusively for primary statistical tests; pseudo-labels (promoted under ≥0.90 probability and positive centroid-cosine) are included only for sensitivity/exploratory analyses.

**Table S1:** Per-gene contingency and label counts used across analyses. For each gene (*GLE1, NUP214, NUP62*), we report curated (ClinVar) missense counts (benign, pathogenic), high-confidence pseudo-labels (probability ≥0.90; benign, pathogenic), and the combined totals. Where applicable, 2 × 2 counts used for Fisher’s exact tests are also shown. Primary statistical claims use the curated set; PU-expanded counts support sensitivity/exploratory analyses.

| Gene | Curated Benign | Curated Pathogen | PU Benign | PU Pathogen | Total Labeled |
| --- | --- | --- | --- | --- | --- |
| *GLE1* | 9 | 6 | 37 | 29 | 81 |
| *NUP214* | 6 | 1 | 16 | 48 | 71 |
| *NUP62* | 7 | 1 | 59 | 66 | 133 |
| **Total** | **22** | **8** | **112** | **143** | **285** |

**Table S2:** Summary of Mann-Whitney U tests across features (pLDDT, BLOSUM62, dHydro, dCharge).

| Feature | U-Statistic | P-value | Δmedian |
| --- | --- | --- | --- |
| dHydro | $U = 246.5$ | $P = 0.02741$ | -3.30 |
| dCharge | $U = 141.0$ | $P = 0.2$ | +0.00 |
| Grantham | - missing | - | - |
| pLDDT | $U = 287.5$ | $P = 0.0002542$ | +59.26 |
| BLOSUM62 | $U = 266.5$ | $P = 0.004409$ | -1.00 |

**Table S3:**
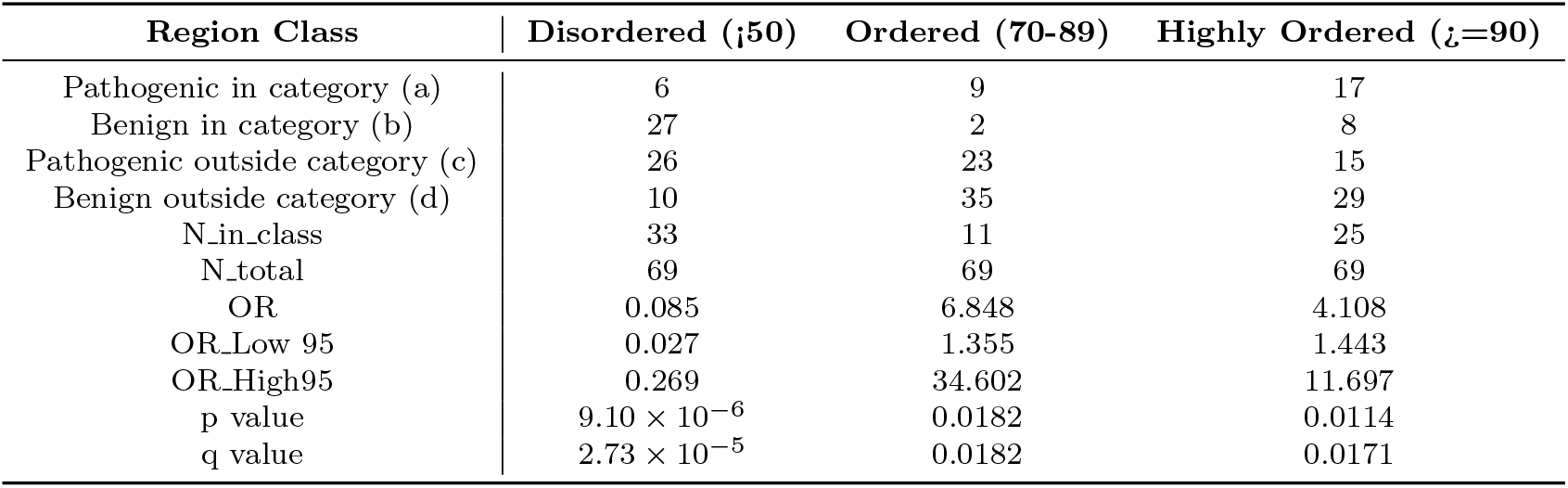
2 × 2 contingency tables and Fisher’s exact test results for enrichment of pathogenic versus benign variants across the indicated categories. Reported statistics include the odds ratio (OR), exact 95% confidence interval (95% CI), the two-sided Fisher p value, and the Benjamini–Hochberg FDR-adjusted q value. OR ¿ 1 indicates enrichment of pathogenic variants within the category; OR ¡ 1 indicates depletion. Unless noted otherwise, results use curated (ClinVar) labels only.

**Table S4:** Mann–Whitney U (two-sided) comparing pathogenic vs. benign in the high-confidence cohort (ClinVar + pseudo *≥* 0.90). Δmedian is median(pathogenic) *™* median(benign).

| Feature | $U$ | $p$-value | Δmedian |
| --- | --- | --- | --- |
| dHydro | 246.5 | 0.02741 | −3.30 |
| dCharge | 141.0 | 0.20000 | +0.00 |
| BLOSUM62 | 266.5 | 0.004409 | −1.00 |
| pLDDT | 287.5 | 0.0002542 | +59.26 |

## References

R. L. Adams and S. R. Wente. Dbp5 associates with rna-bound ip6-gle1 to modulate mrnp remodeling and export. RNA Biology, 14:1–9, 2017.

L. Basel-Vanagaite et al. Mutations in nup62 cause infantile bilateral striatal necrosis. Annals of Neurology, 60:214–222, 2006.

Martin Beck and Ed Hurt. The nuclear pore complex: understanding macromolecular transport through structure, 2017.

Jessa Bekker and Jesse Davis. Learning from positive and unlabeled data: A survey. Machine Learning, 109:719–760, 2020. doi: 10.1007/s10994-020-05877-5.

Yoav Benjamini and Yosef Hochberg. Controlling the false discovery rate: A practical and powerful approach to multiple testing. Journal of the Royal Statistical Society. Series B, 57:289–300, 1995.

The UniProt Consortium. Uniprot: the universal protein knowledgebase in 2023. Nucleic Acids Research, 51(D1):D523–D531, 2023. doi: 10.1093/nar/gkac1052.

Charles Elkan and Keith Noto. Learning classifiers from only positive and unlabeled data. In Proceedings of the 14th ACM SIGKDD International Conference on Knowledge Discovery and Data Mining, pages 213–220, 2008. doi: 10.1145/1401890.1401920.

M. Fornerod, M. Ohno, M. Yoshida, and I. W. Mattaj. Crm1 is an export receptor for leucine-rich nuclear export signals. Cell, 90:1051–1060, 1997. Interacts with CAN/Nup214 at the cytoplasmic face.

Steffen Frey and Dirk Görlich. Fg-rich repeats of nuclear pore proteins form a three-dimensional selective phase. PNAS, 104:11591–11596, 2007.

Björn Hampoelz, Alicia Andres-Pons, Panagiotis Kastritis, and Martin Beck. Structure and assembly of the nuclear pore complex, 2019.

Steven Henikoff and Jorja G. Henikoff. Amino acid substitution matrices from protein blocks. Proceedings of the National Academy of Sciences, 89(22): 10915–10919, 1992. doi: 10.1073/pnas.89.22.10915.

L.-E. Jao et al. Gle1 variants cause a spectrum of motor neuron disease phenotypes. American Journal of Human Genetics, 100:562–572, 2017.

John Jumper, Richard Evans, Alexander Pritzel, and et al. Highly accurate protein structure prediction with alphafold. Nature, 596:583–589, 2021.

Konrad J. Karczewski, Laurent C. Francioli, Grace Tiao, Beryl B. Cummings, Jessica Alföldi, Qiaolin Wang, Ryan L. Collins, Kristen M. Laricchia, Andrea Ganna, Daniel P. Birnbaum, et al. The mutational constraint spectrum quantified from variation in 141,456 humans. Nature, 581(7809):434–443, 2020. doi: 10.1038/s41586-020-2308-7.

Kevin E. Knockenhauer and Thomas U. Schwartz. The nuclear pore complex as a flexible and dynamic gate, 2016.

Melissa J. Landrum, Jennifer M. Lee, Mark Benson, Garth R. Brown, Chen Chao, Shanmuga Chitipiralla, Baoshan Gu, Kenitin Hart, Douglas Hoffman, Wonhee Jang, Smriti Karapetyan, Aleksandr Katz, Justin Liu, Kartik Ramaswamy, Manan Riley, et al. ClinVar: improving access to variant interpretations and supporting evidence. Nucleic Acids Research, 46(D1):D1062–D1067, 2018. doi: 10.1093/nar/gkx1153.

Jaina Mistry et al. Pfam: The protein families database in 2021. Nucleic Acids Research, 49(D1): D412–D419, 2021. doi: 10.1093/nar/gkaa913.

Valeria Nofrini, Daniela Di Giacomo, and Cristina Mecucci. Nucleoporin genes in human diseases, 2016.

H. O. Nousiainen et al. Mutations in gle1 cause lethal congenital contracture syndrome 1. Nature Genetics, 40:155–157, 2008.

Michael P. Rout and John D. Aitchison. The nuclear pore complex as a transport machine. Journal of Biological Chemistry, 278:12521–12524, 2003.

H. B. Schmidt and D. Görlich. Nup98 fg domains from diverse species spontaneously phase-separate into particles with nuclear pore-like permselectivity. eLife, 4:e04251, 2015.

H. E. Shamseldin et al. Nup62-related early-onset neurodegeneration expands the spectrum of nuclear pore complex disorders. Brain, 140:e56, 2017.

Mihaly Varadi, Sarah Anyango, Mandar Deshpande, and et al. Alphafold protein structure database: massively expanding the structural coverage of protein-sequence space with high-accuracy models. Nucleic Acids Research, 50:D439–D444, 2022.

Catherine S. Weirich, Jan P. Erzberger, Keir Flick, et al. Activation of the dead-box protein dbp5 by gle1 and inositol hexakisphosphate is required for mrna export. Nature, 444:351–355, 2006.

